# IraM Remodels the RssB Segmented Helical Linker to Stabilize σ^s^ against Degradation by ClpXP

**DOI:** 10.1101/2023.01.07.523045

**Authors:** C Brugger, S Srirangam, Alexandra M Deaconescu

## Abstract

Upon Mg^2+^ starvation, a condition often associated with virulence, enterobacteria inhibit the ClpXP-dependent proteolysis of the master transcriptional regulator, σ^s^, via IraM, a poorly understood anti-adaptor that prevents RssB-dependent loading of σ^s^ onto ClpXP. This inhibition results in σ^s^ accumulation, and expression of stress resistance genes. Here we report on the structural analysis of RssB bound to IraM, which reveals that IraM induces two folding transitions within RssB, which are amplified via a segmental helical linker. This work highlights the remarkable structural plasticity of RssB and reveals how a stress-specific RssB antagonist modulates a core stress response pathway that could be leveraged to control biofilm formation, virulence, and the development of antibiotic resistance.

## INTRODUCTION

Regulated proteolysis is a major mode of tuning biological activities in all domains of life. In bacteria, it is carried out by ATP-dependent proteolytic machines such as ClpXP. ClpXP is composed of an unfoldase – ATP-dependent hexameric ClpX, and of a sequestered fourteen-subunit peptidase – ClpP, which collaborate to unfold, translocate and then degrade substrates of remarkably diversity^1^. Prime targets of ATP-dependent proteolysis are sigma factors such as *Streptococcus mutans* σ^X 2^, *Streptomyces albus* AntA^3^, *Escherichia coli* σ^32 4^, and especially *Escherichia coli* σ^s^, which was the first sigma factor to be identified as a ClpXP substrate^5-9^. These proteins play important roles in bacterial physiology as they serve as alternative promoter specificity subunits of RNA polymerase that effect wholesale shifts of gene expression profiles in response to developmental or environmental cues^10^. Thus, sigma factors are now widely appreciated as playing key roles in development, motility and chemotaxis, biofilm formation, virulence, and more generally in adaptation to stress^11^. Given that proteolysis disposes of factors irreversibly, correct substrate choice by proteolytic systems is essential, and is tightly regulated spatiotemporally by adaptors, factors that bind a single or small set of substrates and deliver them to the protease^12^.

σ^s^, the master regulator of the *E. coli* general stress response, has a dedicated ClpXP adaptor, a response regulator called RssB^6,7,9,13,14^. RssB has exquisite specificity, and it is the only known σ^s^ adaptor^9^. It acts catalytically under low ratios of RssB: σ^s^ and facilitates the rate-limiting and most poorly understood step in σ^s^ degradation, substrate loading^5,15^. RssB is expressed both in the logarithmic and stationary phase and is subject to homeostatic feedback control via its σ^s^-dependent promoter^16^. The default state of an actively growing, non-stressed cell is to efficiently degrade σ^s 17^. Upon starvation and entry into the stationary phase cells tune down proteolysis of σ^s^, which results in a ∼10-fold increase of the σ^s^ half-life^5^. In fact, upon starvation, bacteria reduce proteolysis of many folded protein substrates, including σ^s 18^. This strategy is beneficial as it ensures speedy return to active growth once the limiting nutrient is reintroduced and is more substrate independent as it relies on the lowering of ATP levels to tune ClpXP activities^18,19^. However, in the case of σ^s^, the dependence on ATP levels is only partial^18^ and ClpXP adaptor antagonists (aka anti-adaptors) provide an alternative strategy for substrate stabilization that is not only adaptor- and substrate-specific, but also dependent on the type of stress encountered. RssB inhibitors (IraD, IraM, IraP) were early identified as ClpXP anti-adaptors via genetic screens^20-24^, and proposed to use distinct mechanisms for RssB inhibition under specific conditions of stress. In the case of IraD, recent structural work has revealed that IraD-bound RssB features a compact architecture with IraD contacting both RssB domains and with RssB featuring a partially disordered segmented helical inter-domain linker (SHL, magenta in Figure 1). Further structural work confirmed that the SHL has increased plasticity that may underlie the rich RssB regulation, both positive (by phosphorylation of the receiver domain, RssB^NTD^)^25,26^ and negative (by stress-specific anti-adaptors that confer some degree of specialization to the general stress response). Some anti-adaptors, such as the IraM paralog IraL, act in the logarithmic phase and have been proposed to contribute to host and niche specificity in uropathogenic *E. coli* strains^27^. Despite their rise to prominence as a novel strategy for transcriptional reprogramming, these inhibitors of RssB activity have remained poorly understood because they are evolutionarily unrelated, and therefore insights gained from the structure of IraD-RssB complexes are not easily generalizable to the other anti-adaptors.

**Figure 1.**
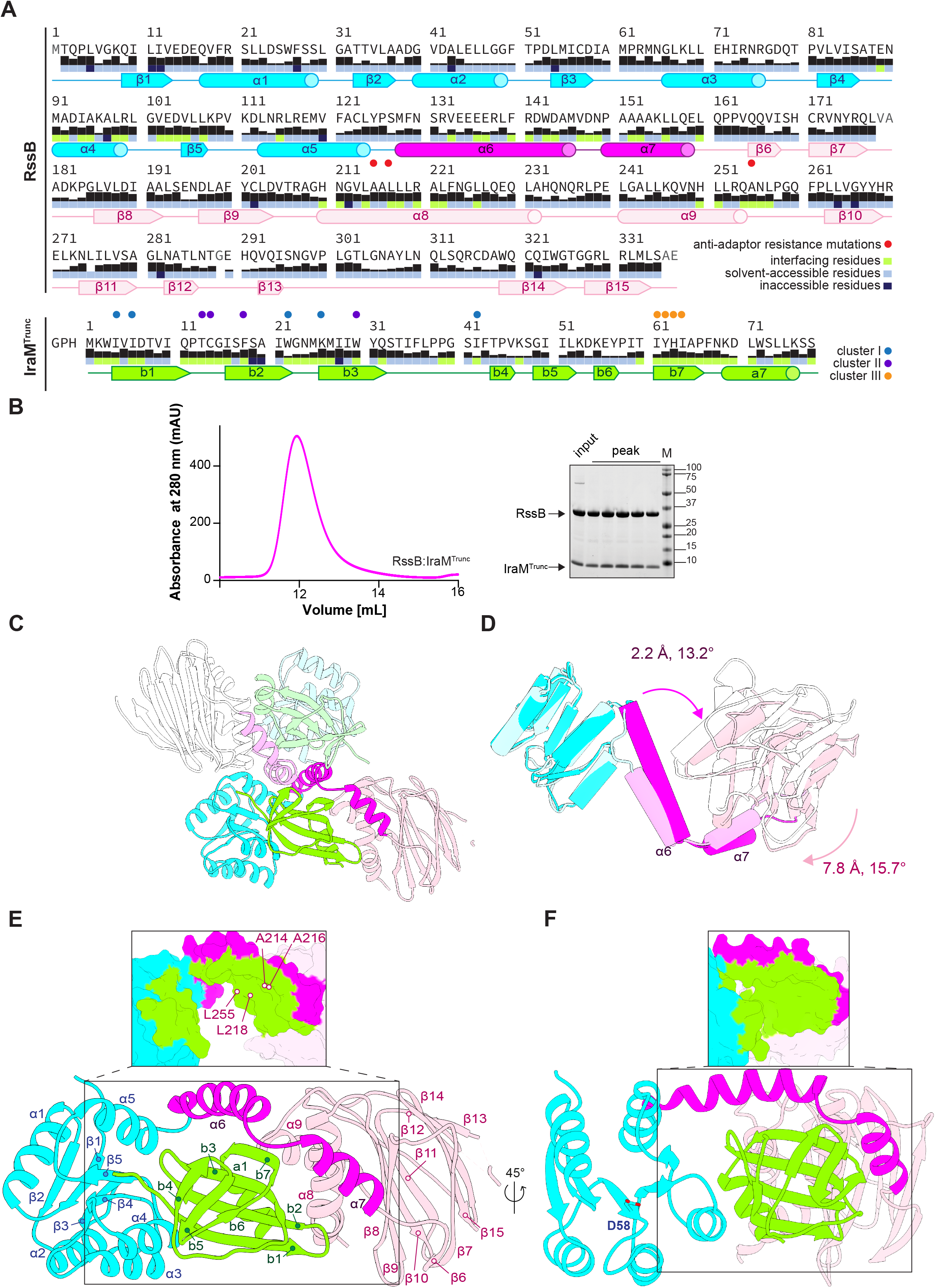
Architecture of the IraM^Trunc^•RssB Complex. A. Schematics of *E. coli* RssB and IraM with relative sequence conservation shown as a black histogram, and residue accesibility indicated by blue (solvent-exposed), navy (buried), and lime (buried at the interface with IraM). Also indicated are residues (red circles) whose substitution leads to resistance to IraM *in vivo* and/or *in vitro* ^20^. B. Size-exclusion chromatography profile of IraM^Trunc^•RssB eluted off a Superdex 75 10/300 column. Also shown is an SDS-PAGE analysis of peak fractions confirming the integrity of the protein complex. C. View of the two copies of the IraM^Trunc^•RssB complex present in the asymmetric unit. Proteins colored by domain as in Figure 1A and with the second copy colored in similar, but more muted hues. D. Overlay of the two RssB copies within the asymmetric unit obtained by a superposition restricted to the α-trace only of RssB^NTD^ (cyan). Note the relative rotation and translation of the SHL and RssB^CTD^. E-F.Front view of *E. coli* IraM^Trunc^ (residues 1 to 78, green) bound to RssB (residues 1 to 337). Insets highlight RssB surfaces buried by IraM binding. Color scheme as in panel A. Lime surfaces indicate RssB:IraM^Trunc^interfaces. The phosphoacceptor site, D58, is highlighted.

To further dissect mechanisms of σ^s^ stabilization, we have isolated *in vivo* assembled complexes of RssB bound to its IraM inhibitor. Anti-adaptor IraM (aka YcgW or ElbA) is induced upon Mg^2+^ starvation^23^, and has also been implicated in resistance to acid, having been described as a “connector” that links activation of the acid-responsive EvgS/EvgA two-component system to the PhoQ/PhoP system^28^, whose activation also controls IraM expression^23^. Critically, both pH shock and Mg^2+^ limitation play important roles in enterobacteria establishing productive infection and virulence. Acidic pH is thought to not only be a stressor, but also a signal that a potential host environment has been reached and thus expression of virulence genes can commence^29^. Mg^2+^ limitation induces expression of virulence factors by flipping the switch from the planktonic to a biofilm lifestyle. In fact, some of the properties characteristic of clinical isolates of *Pseudomonas aeruginosa* can be induced by growth under Mg^2+^ limitation^30^, and the importance of Mg^2+^ homeostasis extends to other bacterial species^31^. At the other end of the spectrum, non-limiting Mg^2+^ dampens starvation-induced persistence^32^, implicating pH sensing, Mg^2+.^ homeostasis, and thus IraM, in the development of antibiotic resistance and expression of virulence genes.

Here we describe a novel conformation of RssB, which we show is induced by IraM binding and results in two folding transitions within the RssB interdomain linker. These changes are amplified by the two helical linker segments and result in a more open RssB conformation, previously proposed to more closely mimic the ON or phosphorylated form of RssB with high affinity for σ^s 25,33^. Surprisingly, we also identify a role for IraM in elevating RssB∼P levels *in vitro*. We propose that RssB inhibition by IraM is due to an effective, tripartite mechanism that leads to 1) occlusion of σ^s^ recognition determinants 2) masking of ClpXP binding motifs and 3) inhibition of RssB recycling by stabilization of RssB∼P. Overall, our data indicate that RssB operates outside of a simple ON (open)/OFF (closed) model of response regulator action.

## RESULTS

### Overall Architecture of the IraM • RssB Complex

We obtained crystals of the *in vivo* assembled IraM•RssB complex and determined its structure using X-ray crystallography (Figure 1 and Table 1). We employed a full-length RssB construct and an IraM variant, IraM^Trunc^ encompassing residues 1-78. IraM^Trunc^ lacks the proteolytically unstable C-terminal region that results in partial cleavage during purification, but, like wild-type, assembles into stable complexes that could be purified to homogeneity (Figure 1B).

**Table 1.**
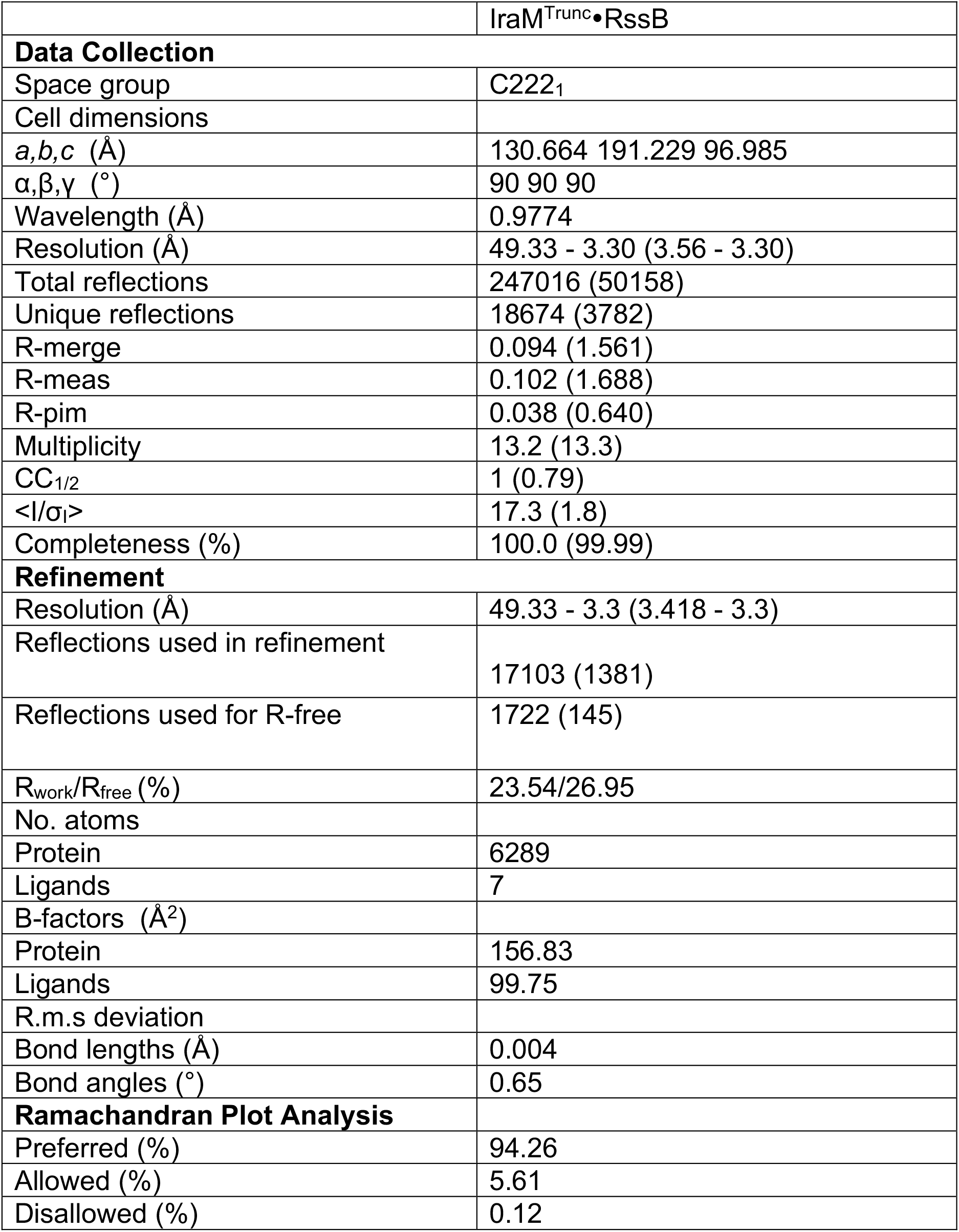
Crystallographic Data and Refinement Statistics. Values in parentheses are for highest resolution shell.

The asymmetric unit of the crystal contains two IraM^Trunc^•RssB complexes (r.m.s.d. of 0.68Å) that dimerize head-to-head via α6 (Figure 1C), part of the SHL of RssB. It has been previously reported that under certain conditions, RssB dimerizes^20,33^. However, structural analysis with PISA^34^ resulted in a low score of 0.1 for the α6− α6 dimerization interface seen *in crystallo*, consistent with the complex being a heterodimer in solution. The two RssB copies in the asymmetric unit are structurally distinct primarily in the SHL, resulting in a small relative rotation of the C-terminal domains in the two RssB copies (Figure 1D).

The IraM core folds into an anti-parallel β−barrel flanked by a β−hairpin on one side and an α−helix on the other (Figure 2A). The C-terminal tail, truncated in our construct, is less conserved and is predicted by AlphaFold2 to be highly flexible^35^. Overall, the IraM fold is unusual and resembles only one other protein model found in the Protein Data Bank, that of the polymyxin resistance protein, PmrD (PDB ID 4HN7, r.m.s.d of 1Å; unpublished), as shown in Figure 2B. In Gram-negative bacteria, two two-component systems are protective and counteract the effect of the last resort antibiotic polymyxin B, PmrA/PmrB and PhoP/PhoQ. The latter pathway is activated upon low extracytoplasmic Mg^2+^ such that the PhoQ histidine sensor kinase phosphorylates PhoP, which promotes transcription of PmrD, which in turn acts as a connector between PmrA/PmrB and the PhoP/PhoQ system. *E. coli* PmrD contains a disulfide bond, but this was not found to be functionally important^36^, and is not present in IraM^Trunc^.

**Figure 2.**
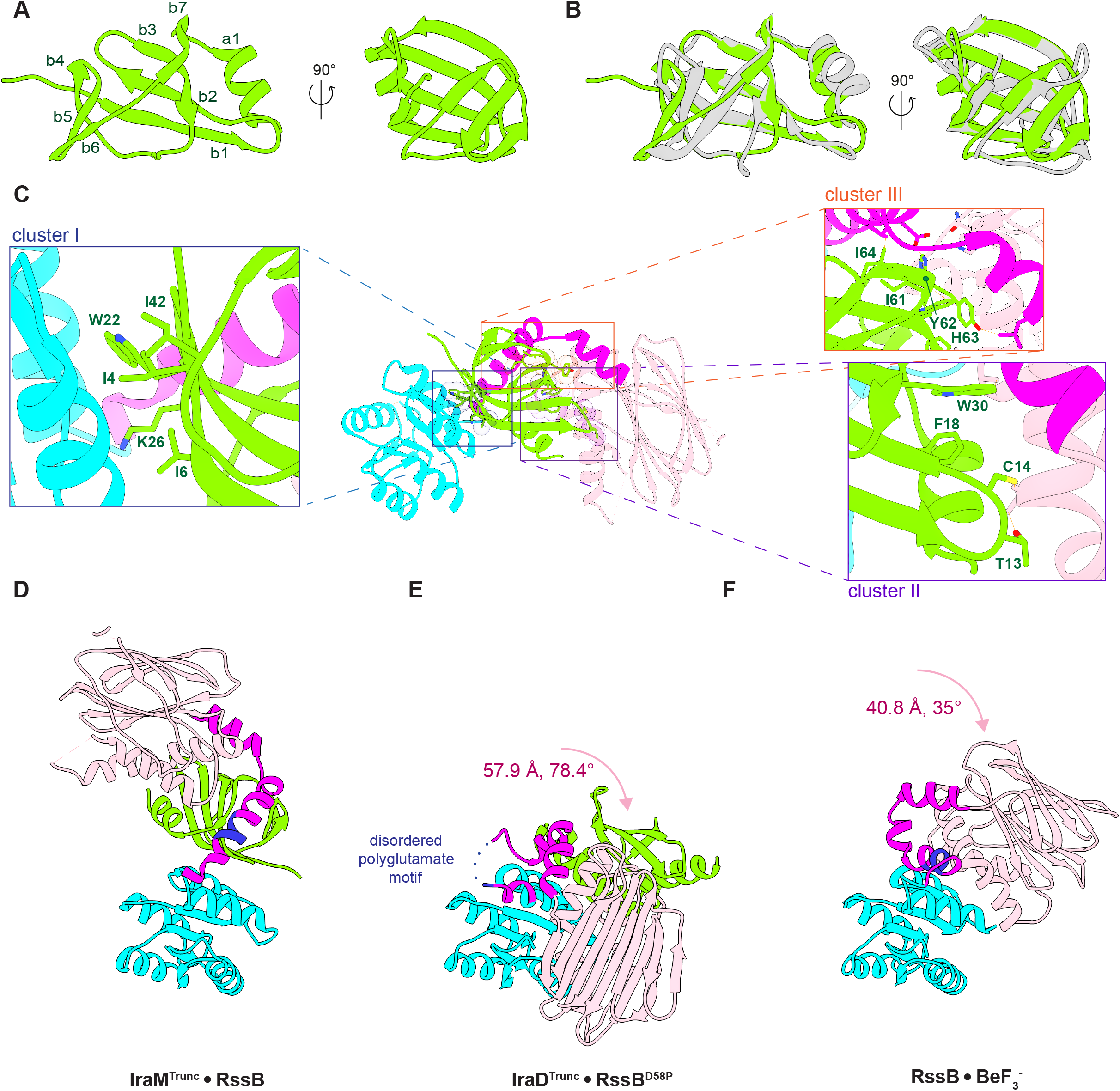
IraM Exhibits a PmrD-like Fold and RssB an Open SHL. A. Views of the IraM fold. B. Superposition between IraM (green) and *E. coli* PmrD (grey). C. View of the IraM^Trunc^•RssB complex colored as in Figure 1A and with insets highlighting the clusters of IraM residues mediating direct contacts with RssB. Selected interface residues are shown as sticks. D-F. Comparison of the RssB conformations observed in the IraM-bound form (D), in the IraD-bound form (E) and bound to the beryllofluoride phosphomimic (F).

The crystallized complex shows a striking disposition of RssB domains, well separated via the SHL, and bridged by one IraM molecule. This abuts both the receiver domain, RssB^NTD^, and the pseudophosphatase domain, RssB^CTD^, and makes extensive interactions with the SHL (Figure 1E,F). Altogether complex formation buries a large ∼3000 Å^2^ of protein surface area, consistent with the stability of the complex. The interface of IraM with RssB is organized around three clusters of IraM residues. Cluster I (Ile4, Ile6, Trp22, Lys26 and Ile42) mediates multiple van der Waals interactions with RssB^NTD^ through its α4-β5-α5 face (Figure 2C, inset). Cluster II (Thr13, Cys14, Phe18 and Trp30 in Figure 2C, inset) interacts with RssB^CTD^ through the signaling helix α8. We note that the IraM•RssB^CTD^ interface (∼ 607 Å^2^, Figure 1E, inset) is larger than IraM•RssB^NTD^ one (∼ 404 Å^2^, Figure 2E-F, insets), consistent with IraM binding to both truncated RssB^CTD^ and RssB^NTD^ *in vitro*^37^ and to RssB^CTD^ only in a bacterial two hybrid assay^20^. This previous work also emphasized the importance of the SHL, which when fused to RssB^NTD^ or RssB^CTD^ enhances the interaction^20^. In fact, residues within strand β7 of IraM (e.g., Ile61, Tyr62, His63 and Ile64, Cluster III in Figure 2C) make multiple van der Waals interactions and are positioned within hydrogen bonding distance from the SHL, from Asp144, Asn149, but also from residues outside of the SHL such as Arg99, His210 and Ala255.

The SHL (magenta in Figures 1-2) adopts a segmented helical structure, akin to what was observed in the IraD^Trunc^•RssB^D58P^ complex^33^ and in beryllofluoride-bound RssB^26^. However, there are key differences in the boundaries of the helical segments due to two folding transitions, one N-terminal to α5, which shortens it by ∼ 1 turn, and one involving the polyglutamate motif in α6 (residues 134-137, Figure 1), which was disordered in the IraD-RssB complex^33^, but here adopts a well-ordered α-helical conformation that results in a straight and extended α6. In contrast, in beryllofluoride-bound RssB, α6 is split into two short α−helical segments (α6a and α6b, Figure 3) to accommodate a large kink. There is currently no structural model available for non-phosphorylated RssB, but IraM can inhibit both RssB and RssB∼P^20^. We note that limited proteolysis in the presence and absence of the small phosphodonor acetyl phosphate failed to reveal conformational differences between RssB and RssB∼P^33^, suggesting that the differences seen here are induced by IraM binding and are not due to the absence of aspartylphosphate.

**Figure 3.**
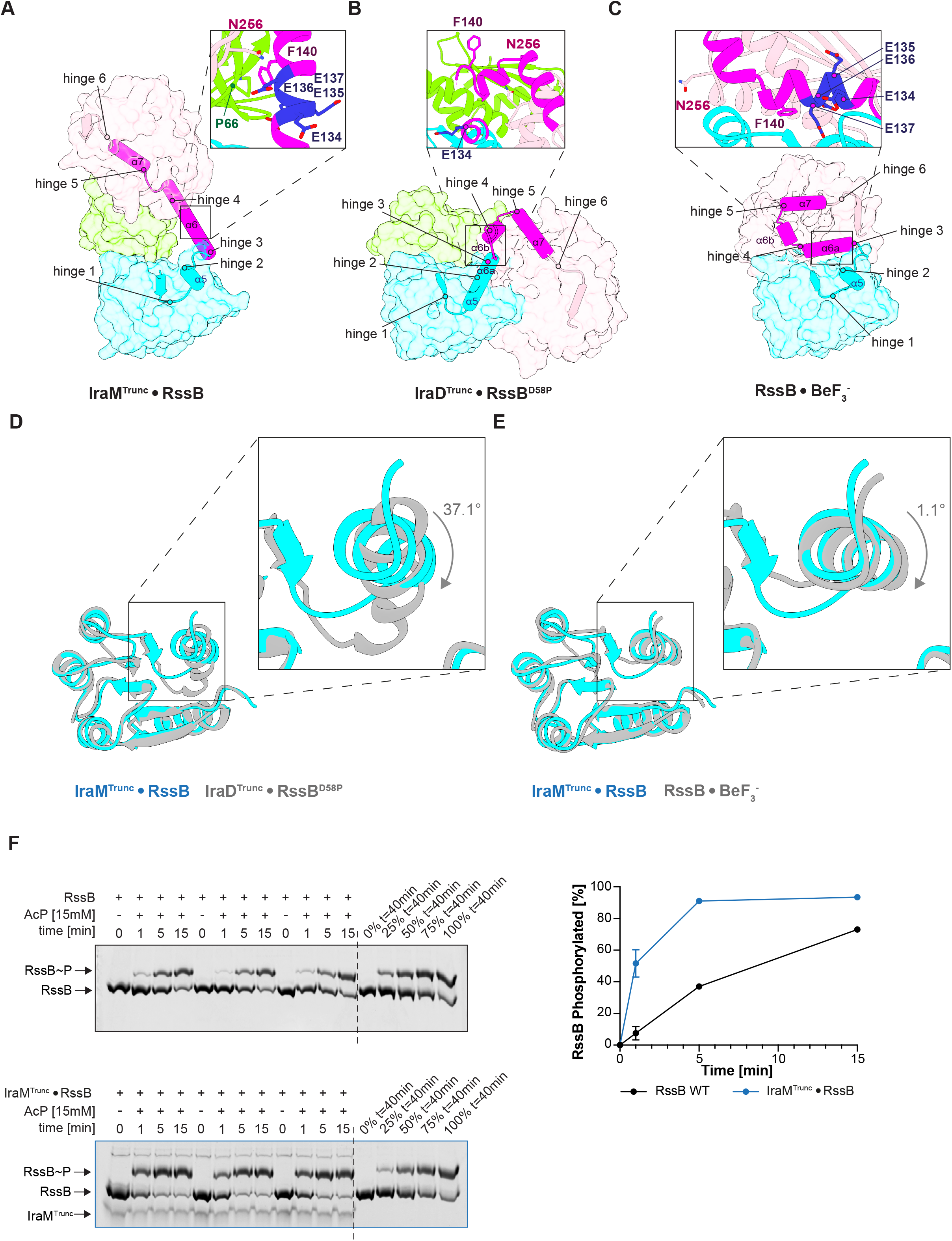
SHL Plasticity. A-C. Views of the SHL in the three known RssB structures: in complex with IraM^Trunc^, with IraD^33^ and bound to beryllofluoride^26^. Shown are putative hinge residues. The polyglutamate motif (E134-E137) is shown in blue and SHL-IraM interactions are highlighted in the insets. D. Superposition between RssB^NTD^ from the IraD^Trunc^•RssB^D58P^ structure (PDB ID 6OD1, grey) and the RssB^NTD^ conformation seen in thee IraM^Trunc^•RssB structure (cyan). E. Superposition between RssB^NTD^ from the beryllofluoride-bound RssB (grey)^26^ and the RssB^NTD^ conformation seen in thee IraM^Trunc^•RssB structure (cyan). Ligands have been omitted for clarity. F. Phos-tag gel analysis of RssB phosphorylation in the absence and presence of IraM and acetyl phosphate. Data obtained for three replicates (left) are plotted on the right as means ± s.d. (n=3). Some symbols are smaller than error bars.

### The Importance of the SHL Hinges for Regulation by IraM

In previous work, we proposed that RssB regulation is controlled mainly by the SHL^33^, which is critical for relaying signals from the phosphoacceptor site, D58, to still poorly-defined sites that associate with α^s^ and/or ClpX^38^. We recently refined this model and proposed that SHL contains hinges that articulate the segments of the linker and are differentially used during regulation^26^. We defined these hinges as follows: Lys108/Pro109 (hinge 1), Leu124 (hinge 2), Phe129/Ser131 (hinge 3), Trp143 (hinge 4), Asn149/Pro150 (hinge 5) and Pro162/Pro163 (hinge 6, Figures 2,3). Hinge Phe129/Ser131 defines the location of a helix-coil transition (compare the linker in the IraD and IraM-bound structures in Figure 3A,B), while Asn149/Pro150 join α6 to α7, and overall result in a structure that is more open compared to the IraD^Trunc^•RssB^D58P^ complex. There, α7 folds back as a rigid body together with RssB^CTD^, leading to extensive RssB^NTD^-RssB^CTD^ contacts via the α1-α5 and α8-α9 faces (Figures 2,3). In contrast to IraD- and beryllofluoride-bound RssB, α6 and α7 are positioned within a crevice and wrap around IraM^Trunc^ and RssB^CTD^.

A region of interest within the SHL is the polyglutamate motif (Figure 3A-C, insets), which was seen to undergo disorder-order transitions in other RssB structures, and here adopts a fully α−helical conformation. Although proximal to IraM, the polyglutamate motif lacks strong contacts with IraM and only makes van der Waals contacts with IraM Pro66, cradled by Glu136, Phe140 but also Asn256 (Figure 3A, inset). While the effect of individual mutations within the polyglutamate motif has yet to be explored, the RssB^AAA^ variant, carrying alanine substitutions of Glu135, Glu136 and Glu137 was proposed to feature a more rigid and perhaps stably helical, open conformation of the SHL^33^. The activity of RssB^AAA^ as an adaptor was significantly compromised suggesting that plasticity in this region may be important for hand-off to ClpXP, yet RssB^AAA^ remains, within the limitations of the *in vivo* assay reported by Dorich et al., sensitive to IraM inhibition^33^. This is consistent with the AAA substitution stabilizing the α−helical form of the polyglutamate motif that supports IraM binding, as seen here.

### IraM and PmrD Both Stabilize the Phosphorylated Form of Their Cognate Receiver

A surprising finding of our study is that IraM-bound RssB^NTD^ more closely resembles the crystallized phosphorylated-like, truncated and beryllofluoride-bound RssB^NTD 25^ rather than the RssB^NTD^ in IraD^Trunc^•RssB^D58P 33^ or the non-phosphorylated truncated, free RssB^NTD^, which was proposed to adopt a meta-active conformation, partially mimicking RssB^NTD^∼P^25^. While the overall differences between IraD^Trunc^ •RssB^NTD^ and IraM^Trunc^•RssB^NTD^ are modest (r.m.s.d. < 1 Å), large differences are seen in the β5-α5 loop and particularly helix α5 (r.m.s.d. of 7.8 Å over the Cα trace of residues 108-132, Figure 3D). In contrast, superposition of beryllofluoride-bound RssB^NTD^ to IraM^Trunc^•RssB^NTD^ results in a low r.m.s.d (∼0.8 Å) even along α5 (Figure 3E). The plasticity of this region of RssB is not unexpected as the 4-5-5 face is one the most dynamic region of receiver domains^39^. However, that IraM stabilizes the RssB∼P like conformation of RssB^NTD^ is surprising given that Battesti et al. presented evidence that IraM prefers RssB or non-phosphorylatable RssB variants, such as RssB^D58P 20^. The corollary of our finding is that IraM may affect RssB∼P levels, by either accelerating the phosphotransfer reaction or decelerating autodephosphorylation or both. Using Phos-tag gel analysis, we therefore probed for RssB phosphorylation in the presence and absence of IraM and AcP. This has been implicated in phosphorylating RssB *in vitro* and *in vivo*^40^. Significantly more RssB∼P can be detected with *in vivo* assembled, purified IraM-RssB complexes than free RssB (Figure 3G). Titrations with purified full-length IraM were not possible due to the pronounced insolubility of the IraM construct. Consistent with our observation, in *Salmonella enterica*, the IraM homolog PmrD functions as a connector between two two-component systems, PmrA/PmrB and PhoQ/PhoP^41,42^ by binding to the phosphorylated PmrA receiver domain and protecting it from dephosphorylation^43^. Thus, IraM shares with PmrD the ability to stabilize the phosphorylated-like conformation of its cognate receiver domain, suggesting that this may be a more broadly applicable property of PmrD-like folds and implicating anti-adaptors in regulating RssB∼P levels, and perhaps RssB recycling.

## DISCUSSION

As a strategy for regulation of gene expression, proteolysis is fast and irreversible since it leads to destruction of the substrate. A correct choice of substrate is therefore essential. In the absence of marking the substrate for destruction such as by ubiquitylation, most bacteria employ adaptor molecules to finely tune the rates of substrate loading and destruction. Adaptors can be broadly classified as “scaffolding” (i.e. functioning as a tether to increase the local concentration of substrate and drive the reaction forward) or “activating”, involving a conformational change within the substrate that makes it engage with the protease, and commit it to degradation. In the case of RssB evidence points to an activating function^5,38,44^, inhibited by multiple, unrelated anti-adaptors, including IraM. Because of the ill-defined nature of the σ^s^ and ClpX binding surfaces and the scarcity of structural information, a unifying model for RssB function and regulation has yet to be achieved. Our study addresses this gap in knowledge and establishes the following.

First, we find that, unexpectedly, IraM stimulates RssB phosphorylation. Ligand stimulation of aspartate phosphorylation has been seen before in both stand-alone^45^ and two-domain response regulators^46,47^ and posits that information transfer between the receiver and the effector domains is bidirectional. In other words, phosphorylation affects the effector domain binding its target, and, in turn, target binding by the effector domain may stimulate phosphorylation of the receiver domain. Invariably, in stand-alone bidirectional regulation was found to involve the “signaling” 4-5-5 face, which is also used by IraM to dock onto RssB^NTD^ (Figures 1,2). A precise mechanistic understanding of this process will require further experimentation on two fronts aimed at (1) clarifying the role played by the SHL hinges and the various interfacial contacts with the 4-5-5-face, and (2) quantifying the rates of the phosphorylation and autodephosphorylation of RssB in the presence/absence of binding partners. However, based on the known activity of PmrD, the closest IraM homolog of known structure, we propose that the main component of IraM-dependent RssB∼P stabilization operates by slowing down autodephosphorylation. Intrinsic dephosphorylation of receiver domains is typically dependent on the nucleophilic attack of a water molecule on the phosphoryl group, which is controlled by the accessibility of this water to the active site^48^. IraM-induced changes within RssB^NTD^ could in principle regulate the transition from active (ON) forms of RssB to inactive (OFF) forms (or vice versa), and directly impact dephosphorylation rates by altering solvent accessibility to the active site. This, however, remains to be ascertained.

In previous work we hypothesized that the SHL serves as the main RssB control element by adopting multiple conformations characterized by different degrees of “openness”^33^. A fully open structure was defined to correspond to the active (ON) form, while “closed” forms were proposed to be OFF, inactive (i.e., with low affinity for σ^s^). We further proposed that all anti-adaptors modulate the affinity of RssB for σ^s^ by interfering with the opening up of the RssB structure^33^. This model was largely based on the compact structure of IraD-bound RssB^D58P 33^ as well as a small number of response regulators that have been crystallized both in their active and inhibited forms^39^. Such work often, but not always, indicated that for activation, tight contacts between the receiver and effector domains must be broken^49,50^. Our present study reveals a surprising, open yet inhibited RssB conformation stabilized by IraM binding, which acts as a RssB^NTD^/ RssB^CTD^ spacer to remodel and open the SHL (Figure 1). While no direct RssB^NTD^/RssB^CTD^ contacts exist, the signaling regions of RssB (the 4-5-5 face and signaling helix α8) are not free but engaged instead in interactions with IraM.

While previous studies together with the data above are consistent with the mutually exclusive nature of IraM and σ^s^ binding^20,26^, they do not completely allow us to delineate all subtleties of the IraM mechanism. Does IraM compete for the same binding site as σ^s^, or does it induce a conformational change that releases σ^s^ from a second site? What drives this conformational change? How does ClpX bind to RssB? Fully answering these questions will require a detailed knowledge of how σ^s^ recognizes RssB upon initial engagement with the protease and upon substrate hand-off. Nevertheless, evidence indicates that both σ^s^ and ClpX have multivalent binding sites that may be used alternatively during engagement and hand-off ^20,26,38^. Data are consistent with σ^s^ binding to RssB^NTD^, perhaps via the 4-5-5 face (where both IraD and IraM dock; both of these anti-adaptors exclude σ^s^)^20^ and perhaps via residues vicinal to the phosphoacceptor site ^51^, but also via unknown determinants within RssB^CTD 38^. Binding to SHL-RssB^CTD^ does not require phosphorylation^38^, while contacts with RssB^NTD^ are directly modulated by phosphoacceptor status^51^. Our structure shows that IraM binding does not block access to the site atop of the Asp58 where σ^s^ was proposed to dock^51^, yet IraM and σ^s^ binding are mutually exclusive^20^, suggesting that other regions in full-length σ^s^ may interact with the 4-5-5 face or that IraM binding exposes determinants that allow ClpX to, at least partially, “eject” σ^s^ by occupying the 4-5-5 face, committing the substrate to degradation and allowing RssB recycling. In fact, using peptide arrays, Micevski et al identified RssB peptides that bind to the zinc-binding domain of ClpX with high affinity and defined a consensus ClpX binding motif, LKxh (where h is a hydrophobic amino acid and x represents any amino acid)^38^. Several such motifs exist in RssB and they span both RssB^NTD^ (LKLL, residues 67-70; LKPV, residues 107-110) and RssB^CTD^ (LKQV, residues 244-248). Strikingly, the motifs within RssB^NTD^ are situated in the immediate vicinity of receiver domain phosphorylation-dependent switches, such as Lys108, substitution of which renders RssB inactive^20^.

While there is ample evidence that phosphorylation via acetyl phosphate increases both the affinity for σ^s^ and degradation rates^5,20,33,38,40^ it is not entirely clear if dephosphorylation is absolutely required for RssB disengaging the protease and subsequent recycling. Thus, at this point in time, we can only speculate that IraM employs a three-pronged mechanistic scheme that inhibits σ^s^ degradation by partially occluding both substrate and protease binding sites (via the 4-5-5 face and helix α9), and also by disfavoring adaptor recycling via stabilizing RssB∼P.

## METHODS

### Protein Expression and Purification

The *E. coli* K12 *iraM* gene was PCR amplified from plasmid pET-IraM ^20^ and subcloned into plasmid pSKB2 (a generous gift from S. Burley) to generate pVF17 (full-length IraM) or pKM6 (IraM^Trunc^). Plasmid pBD1 encoding for RssB was obtained in a previous study ^33^. Resulting plasmids were verified using Sanger DNA sequencing.

For protein co-expression, both IraM^Trunc^ and RssB plasmids were co-transformed into BL21 (DE3) competent cells and grown in Lennox broth (LB) to an OD_600_ of 0.6-0.8 in the presence of 50 *μ*g/mL carbenicillin and 25 *μ*g/mL kanamycin. Cultures were induced with 1mM isopropyl β-D-1-thiogalactopyranoside (IPTG) and grown for 4h at 30°C in a shaking incubator. Hexahistidine-tagged IraM^Trunc^•RssB was purified by Ni^2+^-affinity chromatography, followed by tag cleavage with Prescision protease, a subtractive Ni^2+^-affinity step and size exclusion chromatography using a Superdex 200 16/600 HiLoad or Superdex S200 10/300 columns (Cytiva).

### Crystallization, Structure Determination, and Refinement

IraM^Trunc^•RssB was crystallized at room temperature using sitting drop vapor diffusion with 0.1M sodium acetate pH 4.5 and 120mM magnesium acetate. Crystals were cryo-protected with reservoir solution supplemented with 35% ethylene glycol and flash-frozen in liquid nitrogen.

Data collection was performed at the Advanced Photon Source. Data were indexed with XDS (Kabsch 2010), reduced and scaled with Pointless and Aimless. Molecular replacement was performed with models of the RssB domains obtained from AlphaFold^35^ using the Phaser module^52^ in the Phenix suite. The placed models were further optimized with regards to geometry based on the collected data and re-fed into AlphaFold Multimer to generate optimized models, as recently described^53^. These optimized models were used for a final round of molecular replacement followed by an Autobuild procedure for improved density modification. The resulting maps allowed for building of missing amino acids. Subsequent refinement was carried out in Phenix using the ML refinement target.

### Phosphorylation Assays

RssB and IraM^Trunc^•RssB samples were dialysed against 20mM Tris-HCl pH 8, 150mM NaCl, 15mM MgCl_2_, 1mM DTT, 10% glycerol. 10uL reactions containing 1.5ug of protein and 15mM acetyl phosphate were set up and stopped with 4uL 4xLDS (0.25M Tris-HCl pH6.8, 8% SDS, 40% glycerol) at indicated timepoints. Controls were prepared by phosphorylation of RssB for 40 minutes and mixing with unphosphoylated samples at proportions of 0/25/50/75/100%. Samples were loaded on 10% SuperSep Phos-tag gels (Fujifilm Wako Chemicals U.S.A. Corporation) without prior heating and run at 150V for 70min in the cold using a Tris-Tricine running buffer (0.1M Tris base, 0.1M Tricine, 0.1% SDS). Gels were fixed (10% acetic acid, 50% methanol) for 10 minutes prior to staining with Coomassie G250. Gels were scanned and analysed using ImageLab (Bio-Rad). A correction factor was calculated for each gel based on a RssB calibration curve and internal controls and applied to correct phosphorylation rates of samples. Data was plotted using GraphPad Prism.

## ACKNOWLEDGEMENTS

A.M.D. thanks the Berkeley Center for Structural Biology Beamline 5.01 (Advanced Light Source) for support. This research is based in part upon work conducted using the Rhode Island NSF/EPSCoR Proteomics and Structural Biology Facilities at Brown University. Work in the laboratory of A.M.D. was funded by R01GM121975 and R35GM144124 from the NIH and a Salomon Research Award from Brown University.

